# Nucleotide-dependent conformational changes direct peptide export by the transporter associated with antigen processing

**DOI:** 10.1101/2025.06.12.659373

**Authors:** James Lee, Victor Manon, Jue Chen

## Abstract

The transporter associated with antigen processing (TAP) is essential for adaptive immunity, delivering peptide antigens from the cytoplasm into the endoplasmic reticulum (ER) for loading onto MHC-I molecules. Previous studies have revealed the mechanism by which TAP selectively binds peptides while allowing for sequence diversity, but how the bound peptides are transported and released into the ER is not yet fully understood. Here, we report cryo-electron microscopy structures of human TAP in multiple functional states along the transport cycle. In the inward-facing conformation, ATP binding strengthens intradomain assembly. The transition to the outward-facing conformation is highly temperature-dependent and leads to a complete reconfiguration of the peptide-binding site, facilitating peptide release. ATP hydrolysis opens the consensus site, and the subsequent separation of the NBDs resets the transport cycle. These findings establish a comprehensive structural framework for understanding the mechanisms of peptide transport, vanadate trapping, and trans-inhibition.

## Introduction

The transporter associated with antigen processing (TAP) is an ATP-binding cassette (ABC) transporter essential for adaptive immunity^1,2^. TAP translocates cytosolic peptides, primarily generated by proteasomal degradation of endogenous or foreign proteins, into the lumen of the endoplasmic reticulum (ER). Once in the ER, these peptides are loaded onto major histocompatibility complex class I (MHC-I) molecules for presentation on the cell surface. This process enables cytotoxic T lymphocytes to detect and eliminate cells expressing non-self or aberrant peptides, such as those derived from viruses or malignancies. Mutations in TAP can result in Bare Lymphocyte Syndrome type I (BLS-I), a rare immunodeficiency characterized by impaired antigen presentation and heightened susceptibility to infections^3^. Additionally, numerous viruses, including herpesviruses and adenoviruses, have evolved various strategies to inhibit TAP function, facilitating immune evasion and chronic infection. Structural and mechanistic understanding of TAP are foundational for investigating its function in immune surveillance and exploring potential therapeutic interventions in diseases where TAP function is compromised.

TAP is a heterodimer composed of two homologous subunits, TAP1 and TAP2^4,5^. Each subunit consists of an N-terminal transmembrane domain (TMD0), which interacts with other ER-resident proteins, including tapasin and ERp57^6–13^, to form the larger MHC-I peptide-loading complex. Beyond the TMD0, each subunit contains a core structure composed of six transmembrane (TM) helices that form the peptide translocation pathway, and a cytosolic nucleotide-binding domain (NBD) responsible for ATP binding and hydrolysis. The core TAP heterodimer, excluding the TMD0 regions, is both necessary and sufficient for peptide translocation across the ER membrane^14,15^. While the translocation pathway of TAP is uniquely structured to bind peptides, its NBDs share significant similarity with other ABC transporters. These domains are characterized by highly conserved sequences, including the Walker A and B motifs, which coordinate ATP; the signature motif, a hallmark of ABC transporters; and the D loop, Q loop, and H motif, each named after their conserved residues— aspartic acid, glutamine, and histidine, respectively^16^.

The mechanism for coupling ATP hydrolysis to substrate translocation is highly conserved in the ABC transporter family^17^.The transport cycle of TAP is hypothesized to alternate between two primary conformational states, each exposing the peptide translocation pathway to one side of the membrane. When the NBDs are separated, the translocation pathway is open to the cytosol (inward-facing). When the NBDs form a closed dimer, the transporter opens toward the ER lumen, forming the outward-facing conformation. Upon NBD dimerization, two distinct ATPase sites are formed. One is a catalytically active consensus site composed of the Walker A and B motifs of TAP2 NBD (NBD2) and the signature motif of TAP1 NBD (NBD1). The other is a degenerate site, formed by the corresponding motifs in the opposite NBD, which binds but does not hydrolyze ATP^18–22^.

The inward-facing state of TAP has been directly observed in high-resolution cryo-electron microscopy (cryo-EM) structures determined in the absence of ATP^23^. These studies also elucidate how TAP recognizes and transports a wide variety of antigens — a remarkable feature of the antigen presentation pathway. Peptides, typically 8–14 residues in length^24–26^, bind to TAP within a large transmembrane cavity formed by the TMDs. Each peptide is anchored primarily through backbone interactions at its terminal ends. By prioritizing interactions with the main-chain atoms of the peptide termini, TAP selects peptides of appropriate length while imposes minimal constraints on their sequence^23^. This mechanism underpins TAP’s critical role in enabling the immune system to detect and respond to a broad range of antigens.

To release these peptides into the ER—some of which bind with high affinity—TAP must transition to an outward-facing state. Structural studies of the homologous transporter TmrAB^27^, analyses of isolated NBD1^22,28^, functional characterization of TAP variants with mutations in the NBDs^29^, and FRET measurements in permeabilized cells^30^ collectively support the concept that this transition is driven by NBD dimerization. In this study, we determined cryo-EM structures of outward-facing TAP in two distinct functional states, revealing conformational changes within the NBD dimer upon ATP hydrolysis. Additionally, we determined the structure of inward-facing TAP with either two ATP molecules or an ATP/ADP combination bound to the separated NBDs. These structures provide snapshots of TAP throughout the entire transport cycle, offering a structural model for how ATP powers unidirectional peptide transport from the cytosol to the ER.

## Results

### ATP binding stabilizes NBD1/TMD interface in the pre-translocation state

To analyze the effect of ATP binding, full-length wild-type (WT) TAP was incubated with 5 mM ATP and 20 uM of the high affinity peptide RRYQKSTEL^31^ at 4 °C for 15 sec and immediately vitrified for cryo-EM analysis. Under this condition, TAP adopts an inward-facing conformation with the peptide bound inside the TM cavity and two ATP molecules attached to either NBD (Figures 1 and S1). We did not observe a conformation with the NBDs dimerized, thus it is likely that we have captured TAP in an ATP-bound, pre-translocation state (Figure 1A).

**Figure 1:**
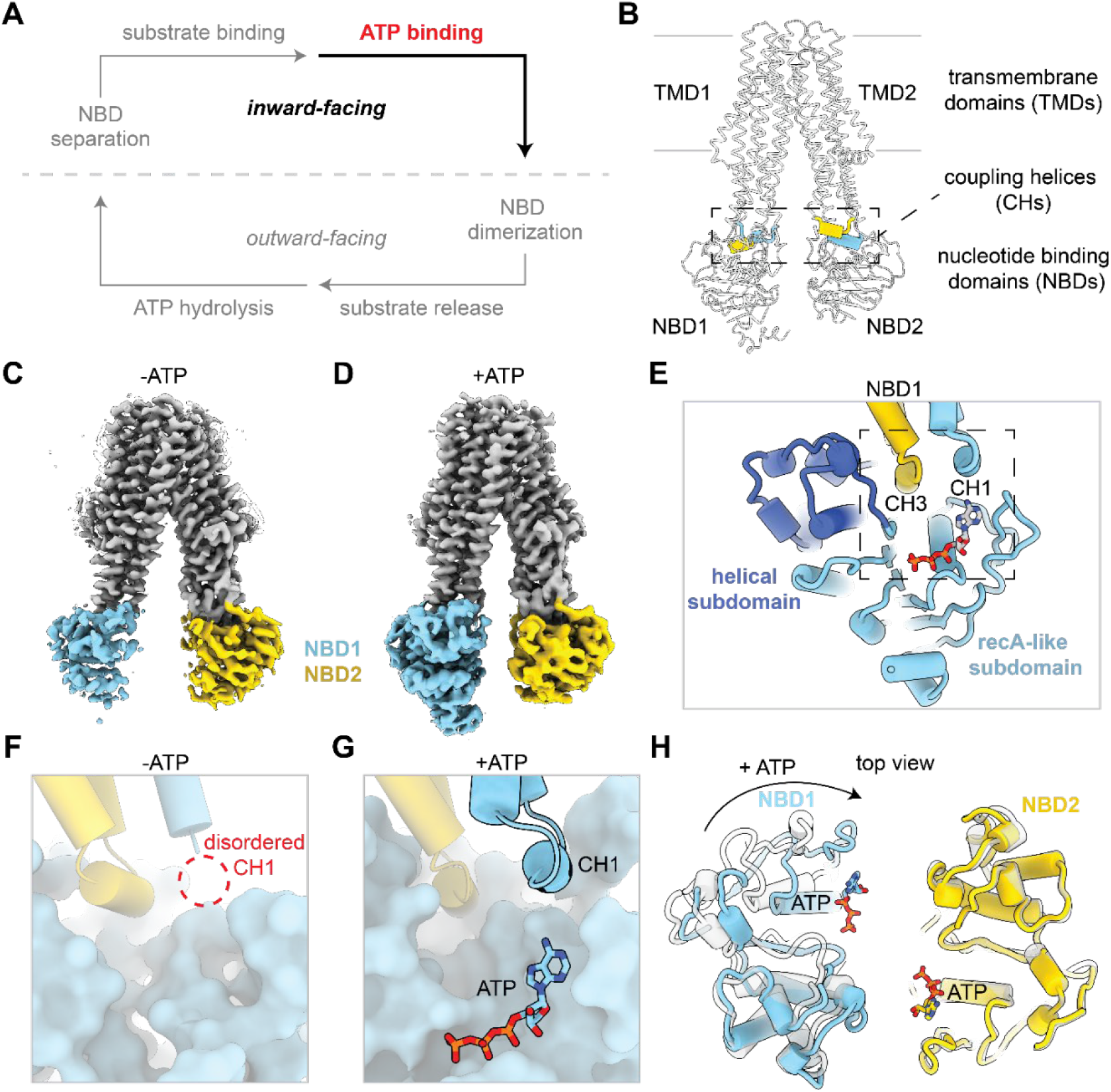
ATP binding stabilizes TAP nucleotide binding domain 1 (NBD1). See also Figure S1. **(A)** Schematic of the TAP transport cycle. ATP binding stabilizes the inward-facing state. **(B)** The coupling helices (CHs) connect the soluble nucleotide binding domains (NBDs) to the transmembrane domains (TMDs). TAP is represented as white ribbon. The TAP CHs are represented as colored tubes. The boundaries of the membrane are shown as silver lines. **(C)** Cryo-EM density of TAP bound to a 9-mer peptide has a flexible NBD1 (EMD-41029). Density corresponding to the TAP TMDs, NBD1, and NBD2 are colored as silver, sky blue, and gold, respectively and contoured to 0.125 standard deviations (SDs). **(D)** Cryo-EM density of TAP bound to a 9-mer peptide and ATP has a rigid NBD1. Density is contoured to 0.8 SDs. **(E)** TAP1 CH1 and TAP2 CH3 interact with NBD1 near the ATP binding site at the interface between TMD and NBD1. TAP is shown as cartoon tubes and ATP is shown as sticks. **(F-G)** Zoomed-in view of TMD/NBD1 interface as boxed in **(E)** in the absence (PDB: 8T4F) **(F)** or presence **(G)** of ATP. The dotted line represents the unresolved CH1. **(H)** ATP binding is associated with a rotation of NBD1 that brings the two NBDs to face each other. Superposition of TAP in the absence (white) and presence (colored) of ATP. The arrow indicates the movement of NBD1 upon ATP binding.

The cryo-EM reconstruction was refined to an overall resolution of 3.6 Å. The densities for the core transporter, peptide, and ATP were well-resolved. However, the densities corresponding to the TMD0 regions of both TAP1 and TAP2 were absent, indicating high flexibility of these regions (Figure S1). Although the overall structure resembles the inward-facing conformation previously observed in the absence of ATP, local differences are apparent in NBD1 and its interface with the TMDs (Figure 1B). In our prior studies, we determined seven cryo-EM structures of inward-facing TAP in the absence of ATP^23^. A common feature of these reconstructions was the poorly defined density for NBD1, which limited us to building only a polyalanine model, even after extensive data processing (Figure 1C). In contrast, the cryo-EM map obtained in the presence of ATP revealed well-resolved density for NBD1, comparable to that of other domains (Figure 1D), allowing us to build its structure *de novo*.

The structure of the NBD and its interface with the TMDs are highly conserved within the ABC transporter family^16,28^. Each NBD comprises two flexibly linked subdomains: a larger RecA-like subdomain and a smaller helical subdomain (Figure 1E). The NBD1/TMD interface involves two intracellular loops: coupling helix 1 (CH1) from TAP1 and CH3 from TAP2, which contact with the Q loop and the helical subdomain in NBD1 (Figure 1E). In the absence of ATP, CH1 lacks defined density, indicating its high flexibility (Figure 1F). In the presence of ATP, CH1 adopts a well-defined structure, forming close contacts with NBD1 and the bound ATP (Figure 1G). Additionally, the linker connecting TMD1 and NBD1 (residues 485–491) also becomes structured in the ATP-bound state, interacting simultaneously with TMD1 and NBD1 to enhance the stability of the NBD1/TMD interface.

A comparison of the inward-facing TAP structures, with and without ATP, shows that the transmembrane domains (TMDs) and NBD2 remain almost identical. However, ATP binding causes the RecA-like subdomain in NBD1 to rotate, moving the degenerate ATP-binding site about 8 Å closer to the molecular center (Figure 1H). As a result, the two NBDs align in the same “head-to-tail” configuration seen in closed dimers. Previously, γ-phosphate-induced rotation of the subdomains was observed in isolated NBD structures^28^. Here, we demonstrate that similar conformational change occurs within the full transporter, suggesting that this rotation is an integral part of the mechanism for coupling transport to ATP hydrolysis.

Together, these results show that ATP binding to inward-facing TAP not only aligns the NBDs structurally but also stabilizes their interactions with the TMD. This process primes TAP for the transition to the NBD-dimerized, outward-facing conformation.

### The outward-facing TMDs exhibit multiple configurations

To stabilize the NBDs in a dimerized conformation (Figure 2A), we replaced the catalytic glutamate in the consensus site with a glutamine, generating a EQ variant that diminishes ATP hydrolysis while preserving ATP binding^32^. The cryo-EM sample was prepared at 4 °C in the presence of 5 mM ATP and in the absence of peptide. Unlike the WT TAP, approximately 50% of the EQ variant adopted the NBD-dimerized conformation (Figure 2B and Figure S2). The overall resolution is ∼3.1 Å; local resolution analysis indicated that the cytosolic half of the molecule is well-defined, whereas the TMDs in the ER leaflet are more mobile (Figure S4). Subsequent focused classification revealed that TAP2 TM3 is conformationally heterogeneous: while the majority of the structures exhibit a continuous helix, a subset forms a kink around P265 near the peptide-binding site (Figure 2C-D and S2-3).

**Figure 2:**
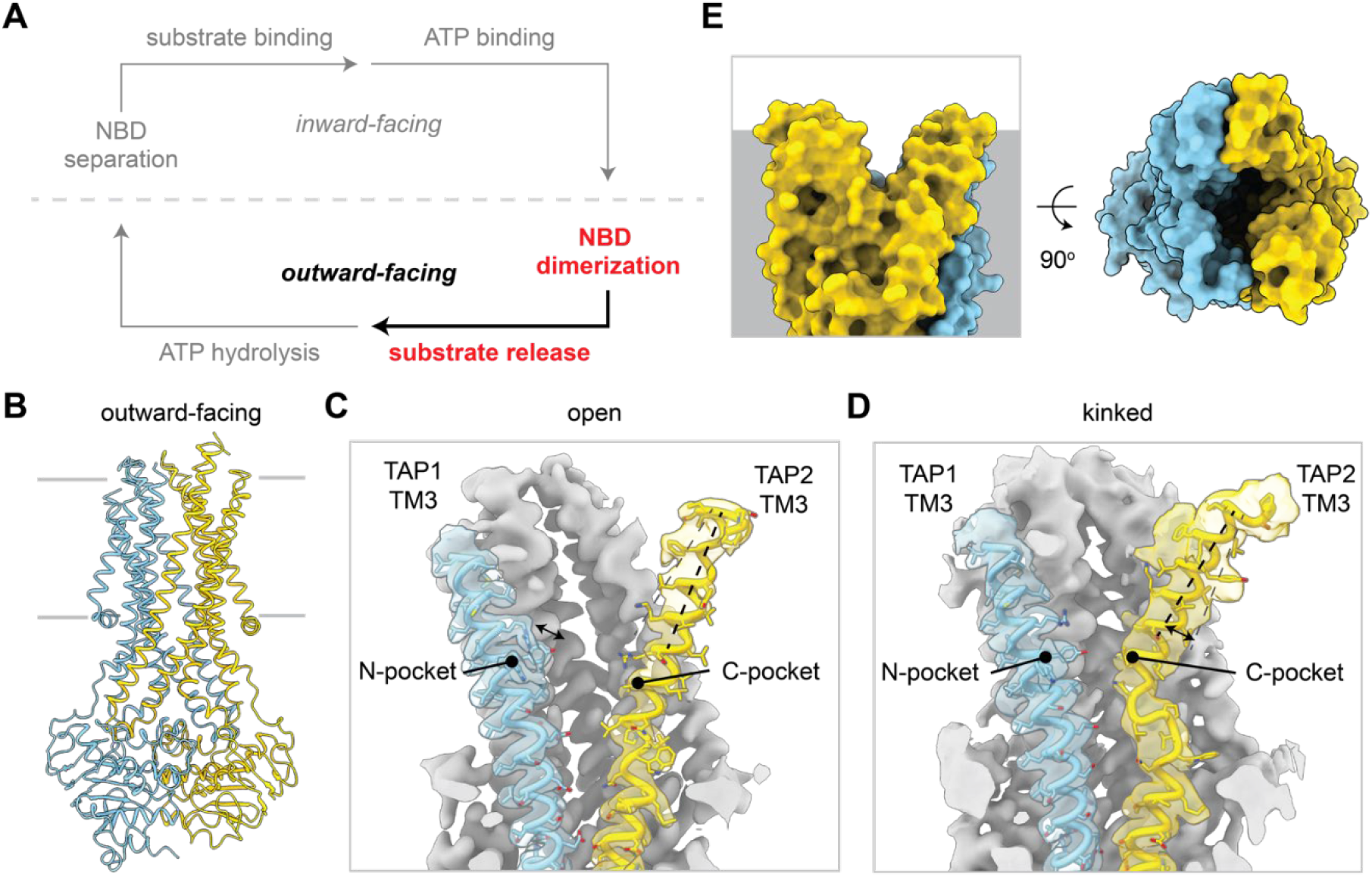
NBD dimerization captures TAP in the outward-facing state. See also Figures S2-S4. **(A)** NBD dimerization stabilizes the outward-facing state. **(B)** Molecular model of TAP (EQ) in the NBD-dimerized outward-facing ATP-bound state. TAP1 and TAP2 are shown as ribbon and colored in sky blue and gold, respectively. The boundaries of the membrane are shown as silver lines. **(C-D)** TAP2 TM3 is flexible and adopts two conformations in the outward-facing state: an outward-facing open **(C)** and kinked **(D)** state. The molecular model of TAP1 and TM3 are represented as cartoons with the side chains represented as sticks. Cryo-EM density corresponding to TAP1 and TAP2 TM3 are colored in transparent sky blue and gold, respectively and contoured to 0.12 and 0.196 SDs, respectively. The N- and C-pockets of the peptide binding sites are labeled. **(E)** The translocation pathway is open to the membrane (top) and endoplasmic reticulum (ER) lumen (bottom). TAP is represented as molecular surface.

The translocation pathway, as anticipated, is open to the ER lumen while sealed off from the cytosol (Figure 2E). Similar to the prototypical bacterial multidrug exporter Sav1866^33^ (Figure S4A), this pathway can also be accessed directly from the lipid bilayer through two lateral openings: one between TM1 and TM2 of TAP1, and another in TAP2 between TM1 and TM3 (Figure S4B). Both openings extend down to the level of the peptide-binding pockets, roughly at the membrane’s midplane (Figure S4B). Compared to the inward-facing conformation, the TM cavity in this state is narrower; and when TM3 adopts a kinked conformation, the translocation pathway becomes even more constricted (Figure 2C-D).

Although the two NBDs are functionally distinct, their structures in this conformation appear very similar, evident by the root mean square deviation (R.M.S.D.) of 0.63 Å over 227 Cα positions (Figure S4C). It is possible that the NBD2 E631Q substitution, by eliminating the catalytic glutamate, effectively converts the consensus site into a degenerate site and thus masks subtle, local asymmetries that may present between the two ATPase sites.

### Transition to the outward-facing state promote substrate release

Transition from the ATP-bound, inward-facing to the outward-facing conformation involves global and local changes (Figure 3). While NBD1 moves as a rigid body (Figure 3A), NBD2 undergoes additional changes in the D loop, which contains a conserved aspartate crucial for NBD dimer stabilization^29^ (Figure 3B). Upon dimerization, D638 in NBD2 shifts 3.6 Å toward the interface, forming two hydrogen bonds with N540 in the NBD1 Walker A motif to further strengthen the NBD dimer interface (Figure 3C).

**Figure 3:**
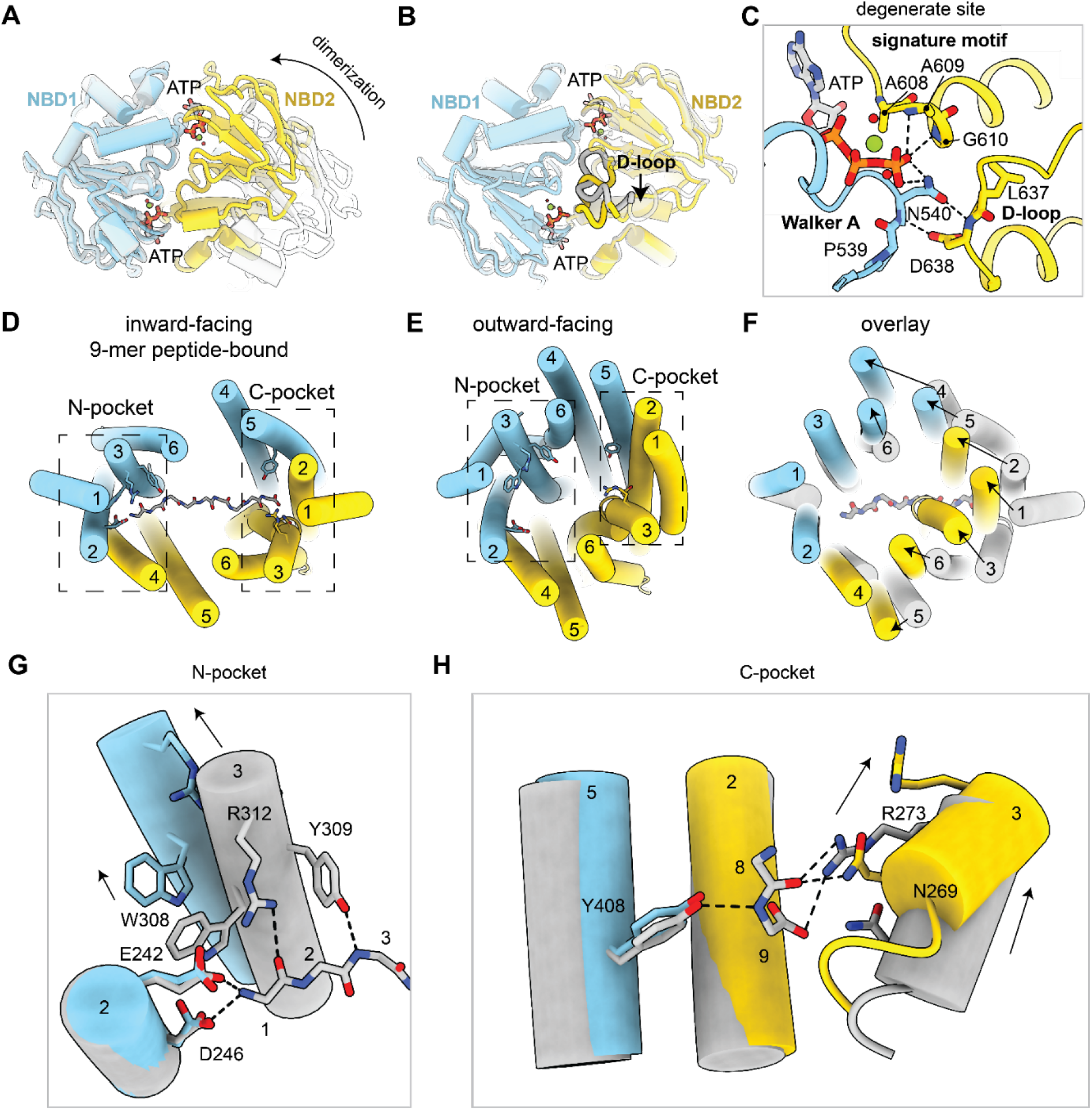
Conformational changes upon NBD dimerization enable substrate release. See also Figures S2-S4. **(A)** Global conformational changes upon NBD dimerization as viewed from the cytoplasm. Superposition of NBD1 in the NBD-separated (white) and NBD-dimerized (colored) conformations. The arrow indicates the movement of NBD2. **(B)** Local conformational changes in the NBDs upon NBD dimerization. Superposition of NBD1 and NBD2 individually in the NBD-separated (white) and NBD-dimerized (colored) conformations. The D-loop is highlighted and the arrow indicates movement of the D-loop to interact with ATP in the degenerate site. **(C)** Zoom-in view of the degenerate nucleotide-binding site in NBD1. Hydrogen bonds are represented as dashed lines. **(D-E)** The TAP translocation pathway in the 9-mer peptide-bound inward-facing **(D)** or outward-facing **(E)** conformation as viewed from the ER lumen. The helices of TAP are numbered and shown as colored cartoon tubes. The main chain backbone of the peptide substrate is shown as silver sticks. The N-pocket and C-pocket are boxed as indicated and residues that comprise each pocket are shown as sticks. **(F)** Superposition of the TAP translocation pathway in the inward-facing peptide-bound (silver) and outward-facing (colored) conformations. Arrows indicate movement of the TAP helices. **(G-H)** Superposition of the N-**(G)** and C-**(H)** pocket of the TAP substrate-binding site as shown in **(D)**. The main chain of the peptide substrate is represented as silver sticks and numbered by alpha-carbon. TAP residues important for interacting with substrate are shown as sticks and hydrogen bonds are shown as dashed lines. Arrows indicate movement of TM3 upon NBD dimerization.

NBD dimerization induces global conformational changes that reorient the translocation pathway, opening it widely toward the ER lumen (Figure 3D-F). These changes are accompanied by a complete reconfiguration of the peptide-binding site. In the NBD-separated conformation TAP binds a peptide through two distal pockets, each interacting with opposite ends of the peptide^23^ (Figure 3D). The N-pocket, primarily composed of charged and aromatic residues in TM2 and TM3 of TAP1, forms extensive contacts with the first three residues of the peptide. Meanwhile, the C-pocket, involving residues mainly in TM 2 and TM3 of TAP2, secures the peptide’s last residue and C-terminus within the translocation pathway. In the NBD-dimerized conformation, the distance between the N- and C-pockets is reduced by approximately 10 Å (Figure 3E), rendering the binding site incompatible with a 9-mer peptide in its extended conformation (Figure 3F). In addition, local structural rearrangements within each binding pocket further reduce peptide affinity. In the N-pocket, TM3 of TAP1 shifts away, disrupting two hydrogen bonds with the peptide (Figure 3G). In the C-pocket, TM3 of TAP2 rotates, moving R273 out of hydrogen-bonding range (Figure 3H). These structural changes not only expose the peptide-binding site to the ER lumen but also reorganize the binding pockets to lower its affinity for peptides, thereby facilitating peptide release.

### Asymmetric opening of the NBDs upon ATP hydrolysis

Since the NBD-dimerized TAP(EQ) structure likely represents the pre-hydrolysis state, where ATP is positioned for hydrolysis and the peptide has already been released to the ER, we aimed to capture the post-hydrolysis state of TAP (Figure 4A). To achieve this, WT TAP was incubated with 10 mM ATP at 37°C for 1 min to facilitate ATP hydrolysis, and the sample was subsequently vitrified for cryo-EM analysis. Under these active turnover conditions, both NBD-separated and dimerized conformations were observed in approximately equal abundance (Figure S5).

**Figure 4:**
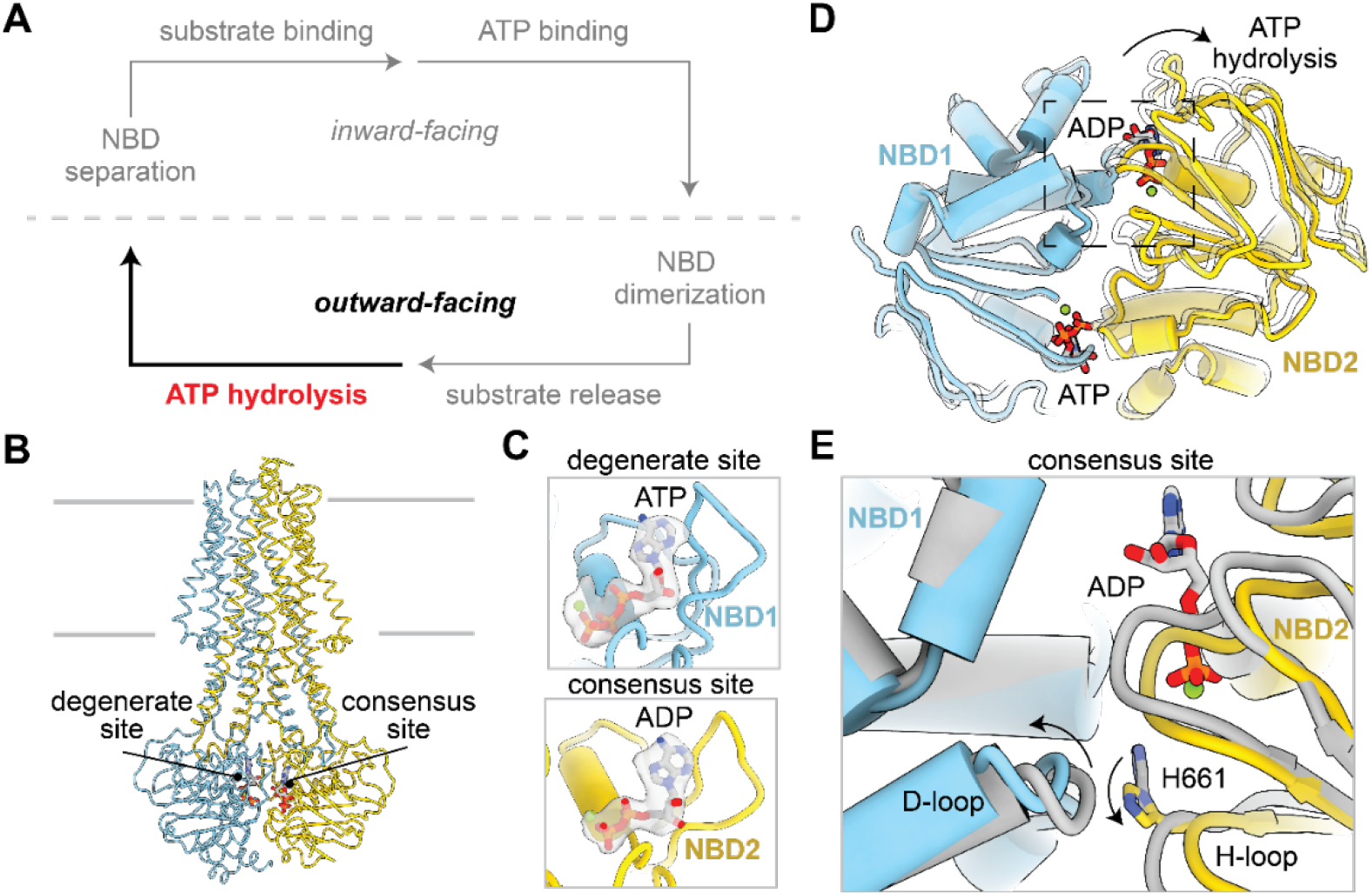
Asymmetric NBD separation after ATP hydrolysis and phosphate release. See also Figure S5. **(A)** The post-hydrolytic state can adopt an outward-facing state. **(B)** Molecular model of wild-type TAP in the NBD-dimerized outward-facing post-hydrolytic state. The degenerate and consensus ATP binding sites are labeled. **(C)** ATP is bound in the NBD1 degenerate site and ADP is bound in the NBD2 consensus site. ATP is shown as sticks and the magnesium atom is shown as spheres. Density corresponding to bound nucleotide is shown as gray surface and contoured to 0.22 SDs. **(D)** Superposition of NBD1 before (white) and after (colored) after ATP hydrolysis as viewed from the cytoplasm. Arrows indicate conformational changes in NBD2 after ATP hydrolysis. **(E)** Zoom-in view of NBD2 consensus site as boxed in **(D)**. Arrows indicate local conformational changes in the D-loop of NBD1 and the H-loop of NBD2 after ATP hydrolysis

The NBD-dimerized reconstruction was refined to 3.2 Å resolution (Figure 4B and S6), showing clear density for ATP at the degenerate site and ADP/Mg^2+^ in the consensus site (Figure 4C), suggesting it represents a post-hydrolytic state before NBD separation. The overall structure closely resembles the pre-hydrolytic, outward-facing conformation (Figure 3), except for a subtle opening at the consensus site (Figure 4D). Following ATP hydrolysis, the H loop of NBD2 (which contains the conserved His residue coordinating the gamma phosphate^34^) and the D loop of NBD1 shift away, resulting in an opening of the dimer interface by approximately ∼3 Å (Figure 4E). No density corresponds to inorganic phosphate (Pi) was observed, suggesting it has already diffused out of the binding site prior to opening of the NBD dimer interface. The asymmetric opening is consistent with the functional asymmetry of the two ATPase site, as ATP hydrolysis only occurs at the consensus site.

The fully separated NBD structure was resolved to a resolution of 3.7 Å (Figure 5A-B and S5), revealing that ATP remains bound at the degenerate site in NBD1 (Figure 5C). The density at the consensus site in NBD2 is weaker and aligns more closely with ADP (Figure 5C). The TMDs have fully reverted to the inward-facing configuration ready to accept substrate (Figure 5A). We interpret this structure as representing a post-hydrolysis state, where the TMDs are poised to recruit a new peptide, and the NBDs are separated to facilitate nucleotide exchange.

**Figure 5:**
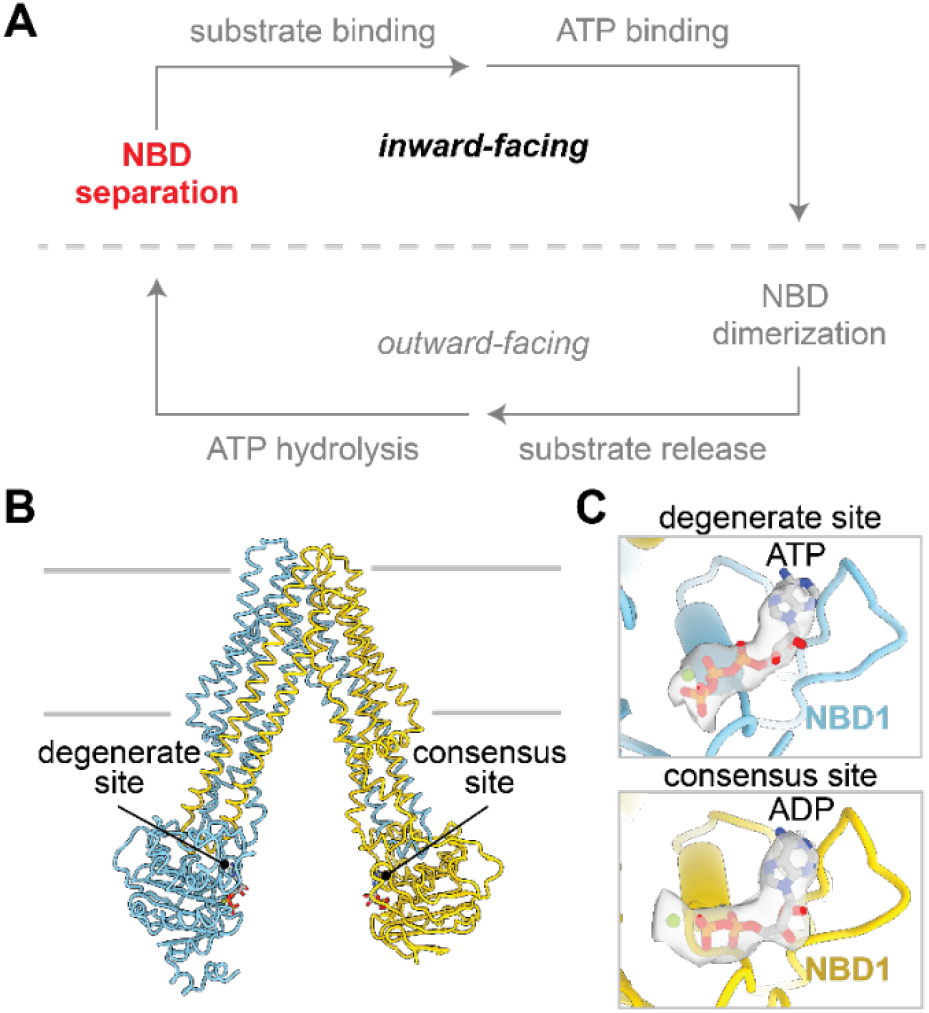
Structure of TAP in the NBD-separated post-hydrolytic conformation. See also Figure S5. **(A)** NBD separation resets the transport cycle. **(B)** Molecular model of wild-type TAP in the NBD-separated outward-facing post-hydrolytic state. Bound nucleotide is shown as sticks and the magnesium atom is shown as spheres. The degenerate and consensus ATP binding sites are labeled. **(C)** ATP is bound in the NBD1 degenerate site (top) while ADP is bound in the NBD2 consensus site (bottom). Density corresponding to bound nucleotide is shown as gray surface and contoured to 0.33 SDs.

## Discussion

This study, in conjunction with a recent publication^23^, has resolved the structure of TAP in six distinct conformational states, allowing us to piece together the complete peptide transport cycle. In the absence of a ligand, TAP adopts an inward-facing conformation with the NBDs widely spaced. Peptide and ATP bind independently^35^, each inducing distinct conformational changes. A peptide of the appropriate length engages both binding pockets within the transmembrane cavity, drawing the TMDs and NBDs closer together. ATP binding stabilizes and strengthens the NBD1/TMD interactions (Figure 1). These coordinated changes drive isomerization to the outward-facing state, where NBD dimerization opens the transmembrane cavity to the ER lumen and reconfigures the binding pockets, reducing their affinity for peptide release (Figure 2 and 3). ATP hydrolysis disrupts the NBD interface (Figure 4), ultimately resulting in complete NBD separation (Figure 5). This mechanism, in which NBD dimerization drives substrate release prior to ATP hydrolysis, is analogous to those observed in the drug transporters Pgp^36^, ABCG2^37^, MRP1^38,39^, and MRP2^40,41^, highlighting a conserved principle within the ABC transporter family.

Consistent with previous FRET studies^30^, we observe that the transition to the outward-facing conformation is highly temperature-dependent. At 4°C, WT TAP predominantly remains in the inward-facing conformation, even in the presence of peptide and ATP. The outward-facing structure was captured either by incubating TAP with ATP at 37°C or by introducing the E631Q substitution to prevent ATP hydrolysis. One possible explanation for the temperature dependence is that higher thermal energy is required to overcome the energy barrier between the inward- and outward-facing conformations. In this scenario, the increased thermal energy allows the NBDs to dimerize with sufficient frequency to be observed by cryo-EM.

For decades, vanadate has been widely used to trap ABC transporters in a conformation presumably mimics the transition state of ATP hydrolysis^42–45^. In the structure of the maltose transporter trapped with vanadate, ADP is tightly bound, with vanadate occupying the position of the gamma-phosphate group of ATP^46^. Vanadate likely binds to an intermediate corresponding to the NBD-cracked-open structure observed in TAP, occurring after Pi has been released but before ADP is discharged and the NBDs fully dissociate (Figure 4). In this conformation, vanadate can access the consensus ATPase site to mimic Pi and stabilize the NBD dimer.

Finally, the structure of the outward-facing TAP also provide a structural basis for understanding trans-inhibition, where peptide transport halts when the luminal peptide concentration reaches 16 µM^29,47^. In this conformation, although both the N- and C-binding pockets are restructured, they likely retain the ability to bind peptides with lower affinity. At micromolar concentrations, peptides may occupy the outward-facing cavity by attaching to one of these pockets, thereby preventing TAP from resetting to its inward-facing conformation.

## Resource availability

### Lead contact

Requests for further information and resources should be directed to and will be fulfilled by the lead contact, Jue Chen.

### Materials availability

All plasmids and reagents generated in this study are available from the lead contact.

### Data and code availability

Cryo-EM density have been deposited in the Electron Microscopy Data Bank under the accession codes EMD-49045, EMD-49046, EMD-49047, EMD-49048, EMD-49049, and EMD-49050. The corresponding atomic models have been deposited in the Protein Data Bank under the accession codes 9N61, 9N62, 9N63, 9N64, 9N65, and 9N66. Any additional data reported in this paper is available from the lead contact upon request.

## Acknowledgements

We thank Rui Yan and Zhiheng Yu at the Howard Hughes Medical Institute (HHMI) Janelia Cryo-EM Facility and Mark Ebrahim, Johanna Sotiris, and Honkit Ng at the Evelyn Gruss Lipper Cryo-EM Resource Center at Rockefeller University for assistance in electron microscopy data collection; and members of the Chen and MacKinnon laboratories for helpful discussions. J.L. is a HHMI Fellow of the Helen Hay Whitney Foundation, and J.C. is an investigator of the HHMI. V. M. is supported by a Medical Scientist Training Program grant from the National Institute of General Medical Sciences of the National Institutes of Health (T32GM007739) to the Weill Cornell/Rockefeller/Sloan Kettering Tri-Institutional MD-PhD Program. This research was supported by the Stavros Niarchos Foundation (SNF) as part of its grant to the SNF Institute for Global Infectious Disease Research at The Rockefeller University.

## Author contributions

J.L., V.M., and J.C. designed experiments; J.L. and V.M. performed experiments; J.L., V.M., and J.C. analyzed data; J.L., and J.C. wrote the paper.

## Supplemental Information

Document S1. Figures S1-S5 and Table S1

### STAR Methods Experimental Model details Cell culture

*Spodoptera frugiperda* Sf9 cells (ATCC CRL-1711) were cultured in Sf-900 II SFM medium (Gibco) supplemented with 5% (v/v) heat-inactivated fetal bovine serum (FBS) (Gibco) and 1% (v/v) antibiotic-antimycotic (Gibco) at 27^°^C. HEK293S GnTI-cells (ATCC CRL-3022) were cultured in Freestyle 293 medium (GIBCO) supplemented with 2% (v/v) FBS at 37^°^C with 8% CO_2_ and 80% humidity. TAP KO cells were generated as previously described. All commercial cell lines were authenticated by their respective suppliers. Commercial cell lines were tested monthly for mycoplasma contamination by PCR using a Universal Mycoplasma Detection Kit (ATCC) and verified to be negative.

### Method details Protein expression

Human TAP constructs were expressed as previously described^23^. Bacmids encoding human TAP fused to a PreScission Protease-cleavable GFP tag were generated in *Escherichia coli* DH10Bac cells (Invitrogen). Baculoviruses were harvested from Sf9 cell media by filtering through a 0.22 µm filter and amplified three times before using for cell transduction. Proteins were expressed in 2L of HEK293S GnTI-cells infected with 5% (v/v) of baculovirus at a density of 2.5-3.0 × 10^6^ cells/ml. Cells were induced with 10 mM sodium butyrate 8-12 hours after infection and cultured at 30^°^C for another 48 hours. Cells were harvested, snap frozen in liquid nitrogen, and stored at -80^°^C.

### Protein purification

Human TAP constructs were purified as previously described. Cells were thawed and resuspended in lysis buffer containing 50 mM HEPES (pH 6.5 with KOH), 400 mM KCl, 2 mM MgCl_2_, 1mM dithiothreitol (DTT), 20% (v/v) glycerol, 1 μg ml^−1^ pepstatin A, 1 μg ml^−1^ leupeptin, 1 μg ml^−1^ aprotinin, 100 μg ml^−1^ soy trypsin inhibitor, 1 mM benzamidine, 1 mM phenylmethylsulfonyl fluoride (PMSF) and 3 µg ml^−1^ DNase I. For samples used for structural analysis, all buffers were supplemented with 1 mM ATP starting from cell lysis. Cells were lysed by three passes through a high-pressure homogenizer at 15,000 psi (Emulsiflex-C3; Avestin). Unbroken cells and cell debris were removed by one low speed spin at 4000g for 15 min at 4^°^C. The supernatant was subjected to a second round of ultracentrifugation at 100,000 x g for 1 hour at 4^°^C in a Type 45Ti rotor (Beckman) to pellet cell membranes. Membranes were resuspended by manual homogenization in a dounce in lysis buffer supplemented with 1% glycol-diosgenin (GDN) (Anatrace) and incubated for 1 hour at 4^°^C. The insoluble fraction was removed by centrifugation at 75,000g for 30 min at 4^°^C and the supernatant was applied to NHS-activated Sepharose 4 Fast Flow resin (GE Healthcare) conjugated with GFP nanobody pre-equilibrated in lysis buffer. After 1 hour, the resin was washed with 10 column volumes of wash buffer containing 50 mM HEPES (pH 6.5 with KOH), 400 mM KCl, 10% glycerol, 1 mM DTT, and 0.01% GDN. To cleave off the GFP tag, PreScission Protease was added to a final concentration of 0.35 mg ml^−1^ and incubated for 12 hours at 4^°^C. The cleaved protein was eluted with 5 column volumes of wash buffer and collected by passing through a Glutathione Sepharose 4B resin (Cytiva) to remove the PreScission Protease. The eluate was then concentrated using a 15 ml Amicon spin concentrator with a 100-kDa molecular weight cutoff membrane (Millipore) and purified by size exclusion chromatography (SEC) using a Superose 6 Increase 10/300 column (GE Healthcare) pre-equilibrated with SEC buffer containing 50 mM HEPES (pH 6.5 with KOH), 200 mM KCl, 1 mM DTT and 0.004% GDN. Peak fractions were pooled using a 4 ml Amicon spin concentrator with a 100-kDa molecular weight cutoff membrane (Millipore) and used immediately for grid preparation or hydrolysis measurements.

### Cryo-EM grid preparation and data acquisition

TAP purified from gel filtration was concentrated to ∼5-6 mg ml^−1^ and, where appropriate, incubated with 150 µM of the corresponding peptide on ice for 30 min. An additional 10 mM of ATP was applied to each sample where appropriate before freezing. To capture the post-hydrolytic state, samples were incubated at 37^°^C for 1 min before freezing. Grids were prepared by applying 3-3.5 µL of sample onto a glow discharged Quantifoil R0.6/1.0 400 mesh holey carbon Au grid with no wait time. The grids were blotted for 3 sec with a blot force of 20 and plunged frozen into liquid ethane using an FEI Mark IV Vitrobot at 6^°^C and 100% humidity.

All cryo-EM data were collected using a 300 kV Titan Krios transmission electron microscope equipped with a Gatan K3 Summit direct electron detector. All micrographs were collected using SerialEM^48^ in super-resolution mode. Data collection parameters for each sample are summarized in Table 1.

### Image Processing

Image processing workflows are summarized in Figures S1, S2, S5, and Table 1. Super-resolution image stacks were gain-normalized, binned by 2, and motion corrected using MotionCor2^49^. Contrast transfer function parameters were estimated using CTFFIND4^50^. Particle picking for all datasets were initially carried out with crYOLO^51^ using its general model, extracted in RELION^52^, and imported into cryoSPARC^53^. The picked particles were subjected to multiple rounds of 2D classification, and the resulting particles were subjected to *ab initio* reconstruction with a maximum resolution set to 9Å. Non-uniform refinement of the best class with the most complete density for the NBDs resulted in a medium-resolution reconstruction of TAP with well-resolved transmembrane helices and NBD2, but with an invisible NBD1. To improve the density of NBD1 in the inward-facing state, all the particles from 2D classification were subjected to iterative rounds of heterogenous refinement using the best reconstruction and a decoy reconstruction with a disordered NBD1 as input models. The resulting particles that gave reconstructions with the most complete NBD1 were then subjected to tandem non-uniform refinement followed by local refinement with a protein mask excluding the micelle.

For TAP(EQ) with ATP dataset, the best particles from these reconstructions were used to train separate Topaz models for both the inward-facing and outward-facing conformations that was used to repick all the micrographs. Particles from both models were combined, duplicates were removed, and subject to the iterative 2D and 3D classification workflow described above. For TAP(EQ) in the outward-facing state, a consensus particle stack was generated consisting of 287,819 particles. This particle stack was imported into Relion using the csparc2star.py script^54^ and subject to Bayesian particle polishing^54^, and refined again in cryoSPARC. Bayesian particle polishing did not improve the other cryo-EM maps.

For TAP(EQ) and ATP in the outward-facing kinked state, the consensus 287,819 particle stack for the outward-facing state was subject to extensive 3D classification in Relion without alignment using a mask excluding the micelle and limiting the resolution E-step to 6Å. Two classes exhibited a difference in the conformation of TAP2 TM3 and were subject to another two rounds of 3D classification in Relion without alignment without limiting the resolution E-step. In both rounds, one class exhibited more continuous density in the TM helices. The final particle stack was then subject to non-uniform refinement to generate the final map.

For the wild-type TAP with ATP at 37^°^C, a combined Topaz model for both the inward-facing and outward-facing conformations was used to pick all the micrographs and subject to iterative 2D classification. *Ab initio* reconstruction with a maximum resolution set to 9Å yielded two distinct classes representing the outward- and inward-facing classes. Each conformation was separately subject to iterative rounds of heterogeneous refinement using the best reconstruction and a lower-resolution decoy reconstruction as input model. Non-uniform refinement of the best class for each conformation was then subject to 3D classification in Relion without alignment using a mask excluding the micelle. For the outward-facing state, the best class was selected based on the completeness of the TMDs. For the inward-facing state, the best stack was selected based on the completeness of the NBDs. The final particle stacks for both conformations were then subject to Bayesian particle polishing and then imported into cryoSPARC for local refinement to generate the final map.

FSC curves were generated in cryoSPARC and resolutions were reported based on the 0.143 criterion. Masking and B-factor sharpening were determined automatically in cryoSPARC during refinement.

### Model building and refinement

The sharpened and unsharpened maps from local refinement were used for model building. Molecular models of TAP under active turnover conditions and of TAP(EQ) with ATP in the inward-facing wide conformation were initially built based on the cryo-EM structure of apo-TAP (PDB: 8T46). The molecular model of TAP with ATP and the b27 peptide was initially built based on the cryo-EM structure of TAP bound to the b27 peptide (PDB: 8T4F). The molecular model of TAP(EQ) with TAP in the outward-facing conformation was initially built based on the cryo-EM structure of TAP bound to the b27 peptide. The initial model was broken into four parts consisting of TAP1 TMD, NBD1, TAP2 TMD, and NBD2. Each model was rigid body fit into the density using ChimeraX^55^ and manually adjusted in Coot^56^. The models were then iteratively edited and refined in Coot, ISOLDE^57^, and PHENIX^58^. The quality of the final models were evaluated by MolProbity^59^. Refinement statistics are summarized in Table 1.

### Figure preparation

Cryo-EM maps and atomic models generated using UCSF ChimeraX^55^. Maps colored by local resolution were generated using cryoSPARC. Structural biology software used in this project was managed by SBGrid^60^. All figures were prepared using Adobe Illustrator.

