## Supplemental Information for "Nucleotide-dependent conformational changes direct peptide export by the transporter associated with antigen processing"

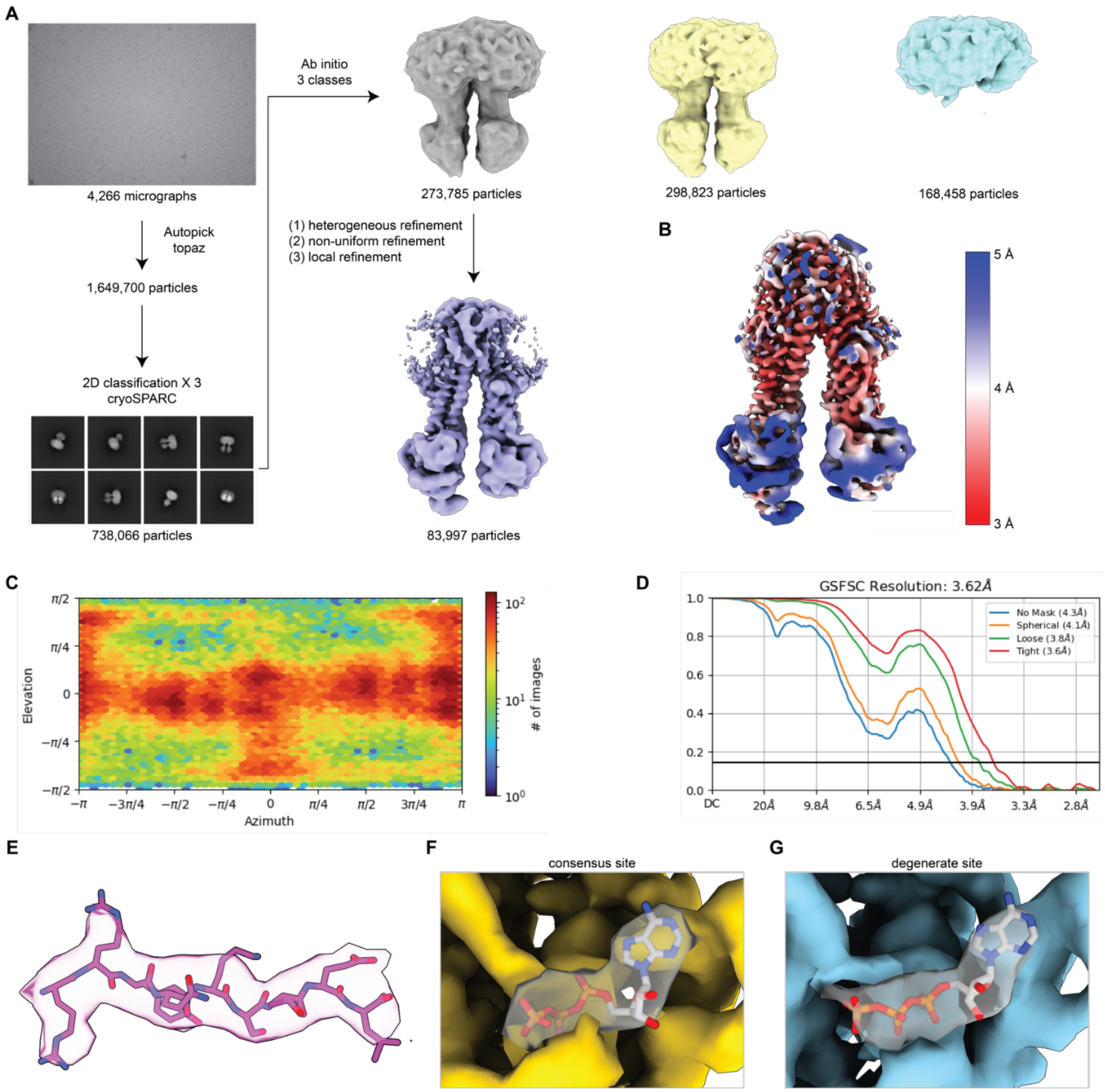


**Figure S1: cryo-EM processing of wild-type TAP in the presence of ATP and the 9-mer peptide RRYQKSTEL at 4ºC. Related to Figure 1.**

**(A)** Summary of image processing. **(B)** Local resolution estimation. **(C)** Angular distribution of particles. **(D)** Fourier shell correlation curve. **(E-G)** Unsharpened cryo-EM density around the peptide (E) and nucleotide binding sites (F-G) contoured to 0.35 and 0.5, respectively.

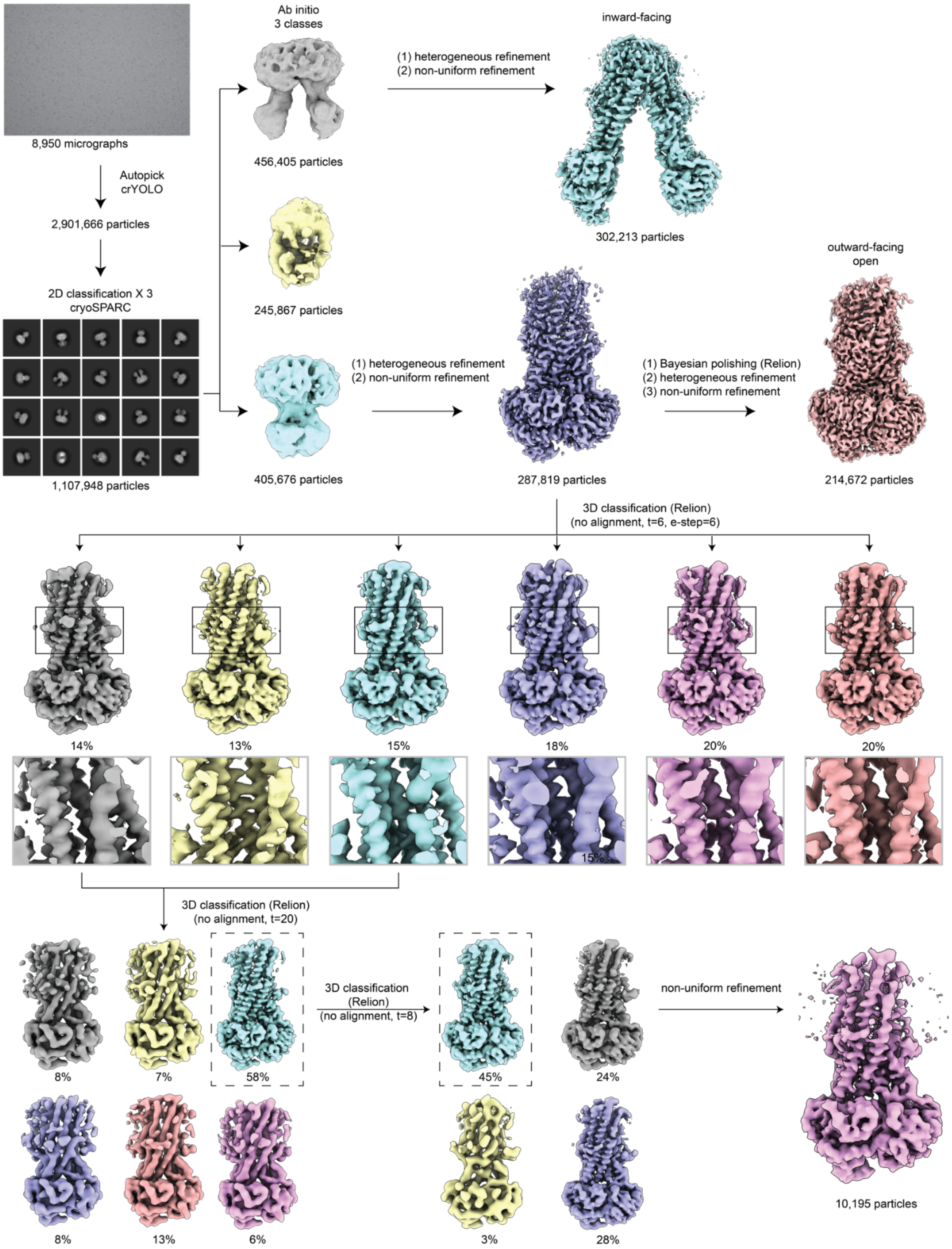


**Figure S2: cryo-EM processing of TAP (EQ) bound to ATP. Related to Figures 2 and 3.**

Summary of image processing.

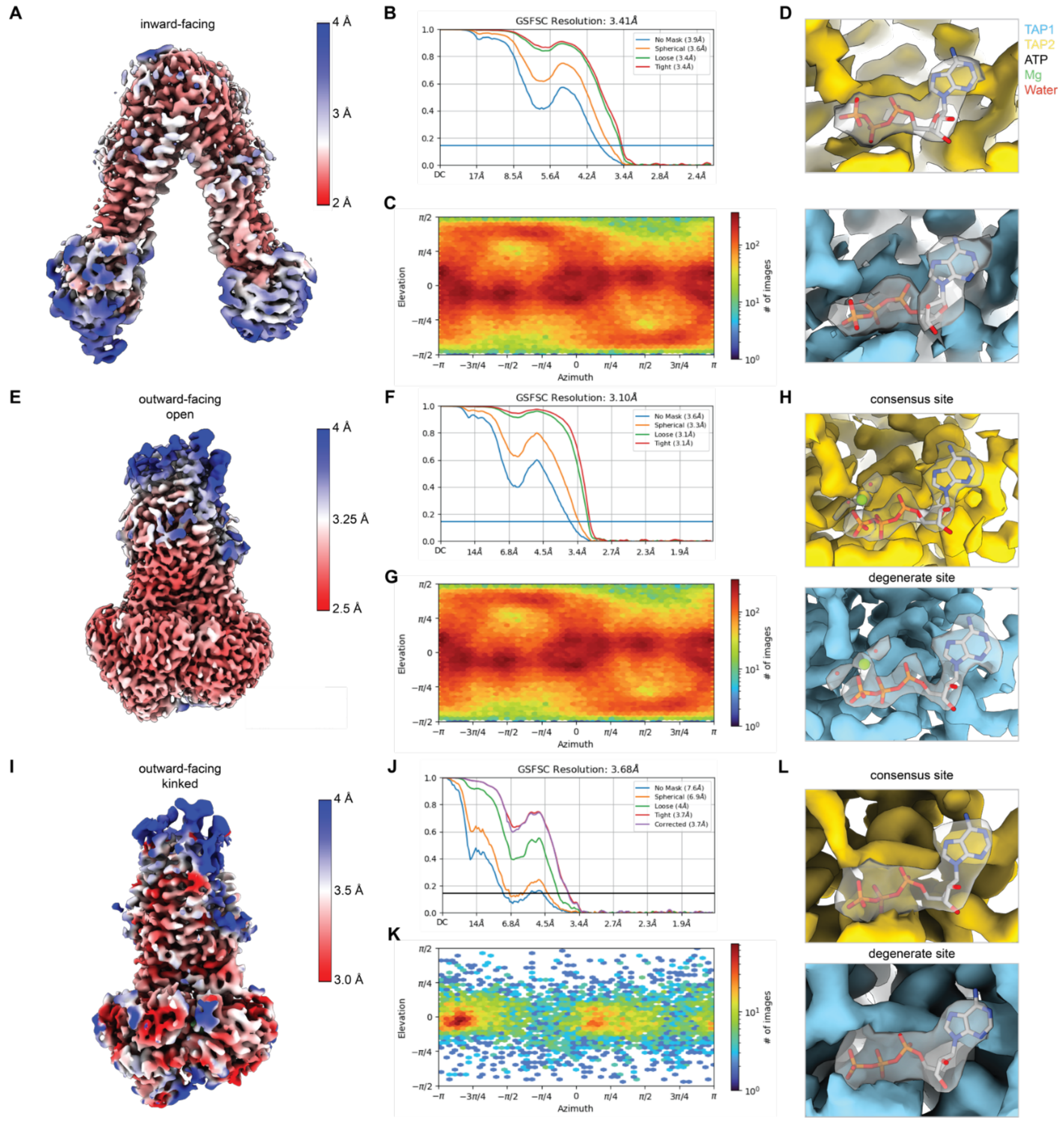


**Figure S3: Cryo-EM analysis of TAP(EQ) in the presence of ATP. Related to Figures 2 and 3.**

**(A-D)** Local resolution estimation **(A)**, Fourier shell correlation curve **(B)**, angular distribution of particles **(C)**, and nucleotide density **(D)** of inward-facing TAP bound to ATP. Reconstructions are contoured to 0.4 SDs. **(E-H)** Local resolution estimation **(E)**, Fourier shell correlation curve **(F)**, angular distribution of particles **(G)**, and nucleotide density **(H)** of outward-facing TAP bound to ATP. Reconstructions are contoured to 0.6 SDs. **(I-L)** Local resolution estimation **(I)**, Fourier shell correlation curve **(J)**, angular distribution of particles **(K)**, and nucleotide density **(L)** of outward-facing TAP bound to ATP. Reconstructions are contoured to 0.4 SDs.

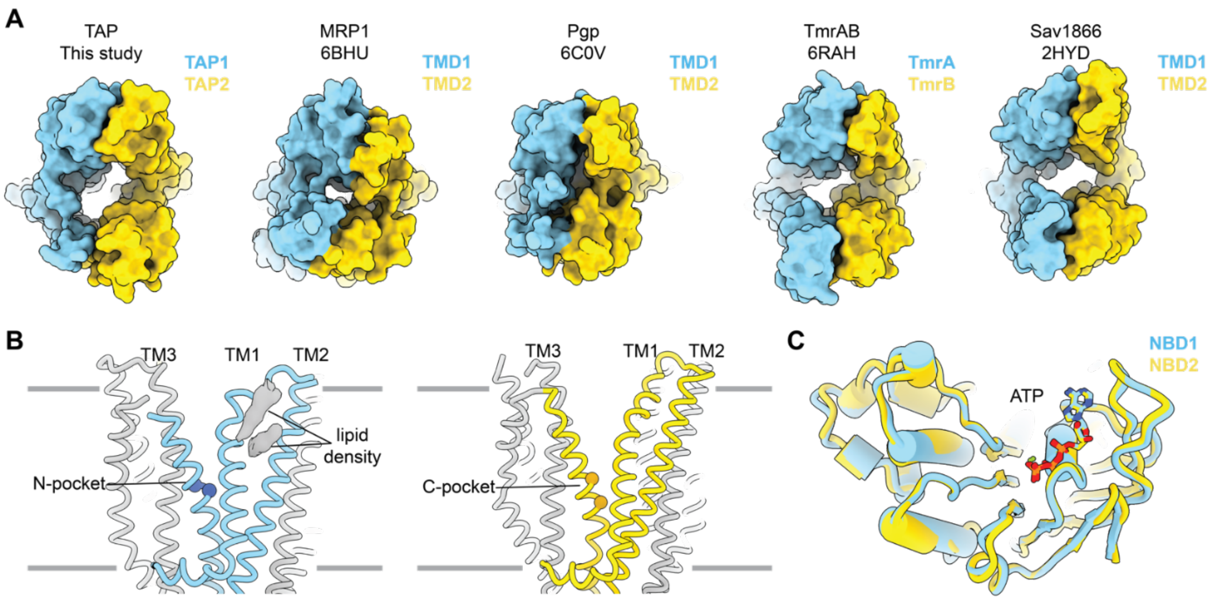


**Figure S4: Structural features of outward-facing TAP(EQ). Related to Figures 2 and 3.**
**(A)** Comparison of outward-facing open TAP to other ABC transporters. Surface representation of various ABC transporters in the outward-facing state as viewed from the ER lumen in the case of TAP or from the extracellular space in the case of the others. Each half of the transporter is colored in sky blue or gold. **(B)** Zoom-in view of the lateral gates of the TAP transmembrane domains from the membrane. TAP1 (left) and TAP2 (right) TM1-3 are colored in sky blue and gold respectively. The alpha carbons corresponding to TAP1 W308 and Y309 in the N-pocket and TAP2 N269k and R273 in the C-pocket are shown as spheres. Density corresponding to putative lipids within the TAP1 lateral gate are marked. Reconstructions are contoured to 0.1. (C) Superposition of NBD1 and NBD2 of TAP(EQ).

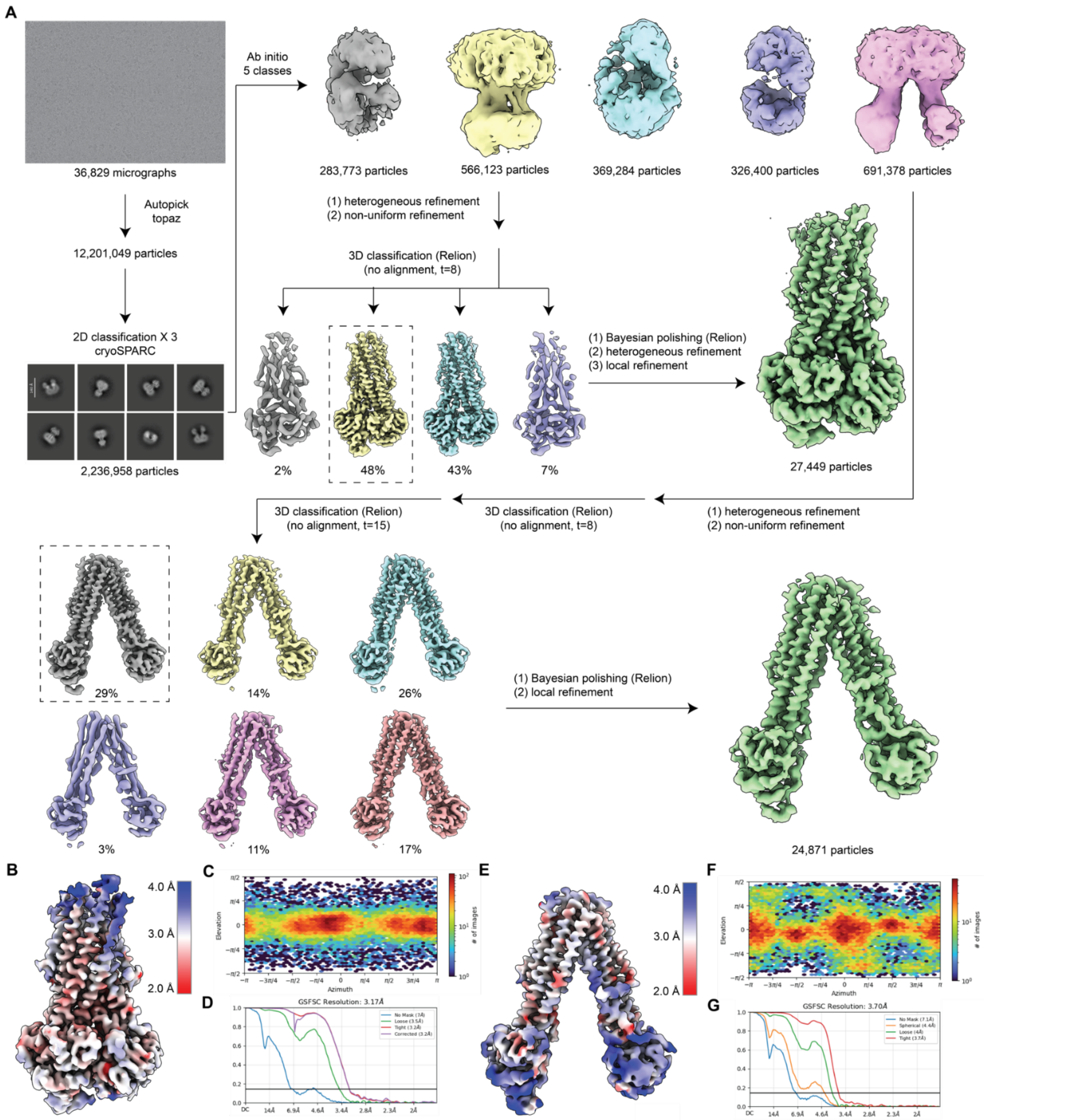


**Figure S5: Cryo-EM analysis of wild-type TAP in the presence of ATP at 37ºC. Related to Figures 4 and 5.**
**(A)** Summary of image processing. **(B-D)** Local resolution estimation **(B)**, Fourier shell correlation curve **(C)**, and angular distribution of particles **(D)** of post-hydrolytic outward-facing TAP bound to ATP/ADP. Reconstructions are contoured to 0.13 SDs. **(E-G)** Local resolution estimation **(E)**, Fourier shell correlation curve **(F)**, and angular distribution of particles **(G)** of post-hydrolytic inward-facing TAP bound to ATP/ADP. Reconstructions are contoured to 0.14 SDs.

**Table S1. Cryo-EM data collection, refinement, and validation statistics**

| | TAP + ATP + B27 peptide Inward-facing EMD-49045 PDB 9N61 | TAP(EQ) + ATP Inward-facing EMD-49046 PDB 9N62 | TAP(EQ) + ATP Outward-facing EMD-49047 PDB 9N63 | TAP(EQ) + ATP Outward-facing Kinked EMD-49048 PDB 9N64 |
|---|---|---|---|---|
| **Data collection and processing** | | | | |
| Magnification | 81,000 | 105,000 | 105,000 | 105,000 |
| Voltage (kV) | 300 | 300 | 300 | 300 |
| Electron exposure ($e^{-}$/Å$^2$) | 50 | 50 | 50 | 50 |
| Defocus range (μm) | 0.8 to 1.5 | 0.8 to 2.0 | 0.8 to 2.0 | 0.8 to 1.5 |
| Pixel size (Å) | 1.09 | 0.847 | 0.847 | 0.847 |
| Symmetry imposed | C1 | C1 | C1 | C1 |
| Initial particle images (no.) | 1,649,700 | 2,900,666 | 2,900,666 | 2,900,666 |
| Final particle images (no.) | 83,997 | 302,213 | 214,672 | 10,195 |
| Map resolution (Å) | 3.6 | 3.4 | 3.1 | 3.7 |
| FSC threshold | 0.143 | 0.143 | 0.143 | 0.143 |
| **Refinement** | | | | |
| Initial model used (PDB code) | 8T4F | 8T46 | 8T4F | 9N63 |
| Model resolution (Å) | 3.8 | 3.5 | 3.1 | 3.9 |
| FSC threshold | 0.5 | 0.5 | 0.5 | 0.5 |
| Map sharpening *B* factor (Å$^2$) | 111.5 | 104.8 | 80 | 68.7 |
| Model composition | | | | |
| Non-hydrogen atoms | 8864 | 8778 | 8583 | 8526 |
| Protein residues | 1130 | 1121 | 1097 | 1089 |
| Water | 0 | 0 | 4 | 4 |
| Ligands | 2 | 2 | 4 | 4 |
| *B* factors (Å$^2$) | | | | |
| Protein | 95.03 | 100.04 | 76.83 | 48.35 |
| Ligand | 139.77 | 129.69 | 46.88 | 37.54 |
| Water | N/A | N/A | 43.90 | 37.35 |
| R.m.s. deviations | | | | |
| Bond lengths (Å) | 0.004 | 0.004 | 0.004 | 0.004 |
| Bond angles (°) | 0.731 | 0.633 | 0.765 | 0.793 |
| Validation | | | | |
| MolProbity score | 1.45 | 1.43 | 1.35 | 1.41 |
| Clashscore | 5.06 | 5.22 | 3.71 | 4.38 |
| Poor rotamers (%) | 0.42 | 0.11 | 0.32 | 0.43 |
| Ramachandran plot | | | | |
| Favored (%) | 96.89 | 97.14 | 96.87 | 96.85 |
| Allowed (%) | 3.11 | 2.86 | 3.13 | 3.15 |
| Disallowed (%) | 0.00 | 0.00 | 0.00 | 0.00 |

**Table S1. Cryo-EM data collection, refinement, and validation statistics**

| | WT TAP + ATP<br><br>Inward-facing<br><br>EMD-49049<br>PDB 9N65 | WT TAP + ATP<br><br>Outward-facing<br><br>EMD-49050<br>PDB 9N66 |
|---|---|---|
| **Data collection and processing** | | |
| Magnification | 105,000 | 105,000 |
| Voltage (kV) | 300 | 300 |
| Electron exposure ($e^{-}$/Å$^2$) | 50 | 50 |
| Defocus range (μm) | 0.8 to 2.0 | 0.8 to 2.0 |
| Pixel size (Å) | 0.86 | 0.86 |
| Symmetry imposed | C1 | C1 |
| Initial particle images (no.) | 5,485,574 | 5,485,574 |
| Final particle images (no.) | 24,871 | 27,449 |
| Map resolution (Å) | 3.7 | 3.2 |
| FSC threshold | 0.143 | 0.143 |
| **Refinement** | | |
| Initial model used (PDB code) | 9N62 | 9N63 |
| Model resolution (Å) | 4.0 | 3.4 |
| FSC threshold | 0.5 | 0.5 |
| Map sharpening *B* factor (Å$^2$) | 93.2 | 53.0 |
| Model composition | | |
| Non-hydrogen atoms | 8778 | 8539 |
| Protein residues | 1121 | 1093 |
| Ligands | 4 | 4 |
| *B* factors (Å$^2$) | | |
| Protein | 49.99 | 57.13 |
| Ligand | 92.44 | 47.33 |
| R.m.s. deviations | | |
| Bond lengths (Å) | 0.004 | 0.003 |
| Bond angles (°) | 0.877 | 0.730 |
| Validation | | |
| MolProbity score | 1.40 | 1.44 |
| Clashscore | 3.74 | 4.37 |
| Poor rotamers (%) | 0.32 | 0.22 |
| Ramachandran plot | | |
| Favored (%) | 96.42 | 96.58 |
| Allowed (%) | 3.58 | 3.42 |
| Disallowed (%) | 0.00 | 0.00 |
